# Human see, human do? Viewing tool pictures evokes tool-use action information in hand-selective occipitotemporal cortex

**DOI:** 10.64898/2026.09.15.751758

**Authors:** Annie Warman, Diana Tonin, Fraser W. Smith, Stéphanie Rossit

## Abstract

Amongst all the objects we encounter, tools are unique because they are tightly linked to predictable actions. Neuroimaging studies have reported selective responses in occipitotemporal and parietal cortices when viewing pictures of tools or hands. Whether these responses contain information about specific tool-use actions (e.g., rotation for keys), and whether such information shares a neural format with action execution, remains unclear. Here, we used fMRI and multivoxel pattern analysis to ask whether specific actions associated with viewed tools could be decoded and whether action-specific information generalised across viewing and acting. Participants (N=18; 11 females) viewed tool pictures and, in separate runs, pantomimed tool-use actions in response to tool names. Familiar tools differed in their associated action (rotate or squeeze) but were matched for grip type. Viewing tool pictures elicited tool-use action information in lateral occipitotemporal cortex (LOTC), parietal cortex, and even somatosensory cortex. Notably, hand-selective LOTC was the only region in which action decoding exceeded tool-identity decoding during both viewing and pantomime, suggesting sensitivity to learned relationships between objects and characteristic hand actions. However, neither region-of-interest nor whole-brain searchlight analyses yielded reliable cross-task decoding. Thus, although overlapping cortical regions contained information about tool-use actions during perception and production, we found no evidence that this information was represented in a shared multivoxel format. These findings suggest that seeing familiar tools evokes distributed information about their associated actions and sensory consequences, but that such information may be expressed in task-dependent formats rather than simply reinstating the neural patterns engaged during action production.

## Introduction

In everyday life, objects provide opportunities for action. Gibson’s (1979) concept of affordances proposed that environmental properties are perceived in relation to the actions they make possible. Tools provide a powerful example because experience tightly associates their visual properties with learned relationships between objects, bodily actions, and their consequences (Johnson-Frey et al., 2005). Thus, we learn not only that keys open doors, but that using them requires a characteristic hand rotation. Recognising a familiar tool may therefore provide information about both what it is and how the body can interact with it to achieve a goal. Understanding these perception-action links is fundamental to explaining skilled object interaction.

Neural models of perception and action propose a left-lateralized tool-processing network involving ventral and dorsal regions (Milner and Goodale, 2006; Rizzolatti and Matelli, 2003; Buxbaum, 2017; Buxbaum and Kalénine, 2010; Lewis, 2006). Ventral regions, including lateral occipitotemporal cortex (LOTC) and posterior fusiform cortex, are implicated in tool perception. Within the dorsal stream, dorsal-dorsal regions, including intraparietal and dorsal premotor cortices, are thought to process structural manipulation knowledge (e.g., how to grasp a tool), whereas ventro-dorsal regions, including middle temporal and supramarginal gyri and ventral premotor cortex, process functional manipulation knowledge (i.e., how to move the hand to use a tool). Grounded theories further propose that conceptual tool representations involve regions associated with both tool perception and use, even when no action is required (Martin, 2016; Barsalou et al., 2003).

Consistent with these accounts, viewing tool pictures elicits sensorimotor activity (Chao and Martin, 2000; Lewis, 2006), interpreted as retrieval of action representations concerning hand movements associated with tool use (Martin et al., 1995; Fang and He, 2005). Viewing hands activates closely overlapping regions, proposed to reflect the shared functional role of hands and tools in skilled object-directed actions (Bracci et al., 2012, 2016; Bracci and Peelen, 2013; Striem-Amit et al., 2017). Left LOTC contributes causally to discriminating whether viewed tools are associated with rotating or squeezing actions (Perini et al., 2014), while hand-selective LOTC contains information about specific hand postures (Bracci et al., 2018). However, whether neural responses to tools contain information about specific associated actions when action is task-irrelevant remains unresolved. Moreover, spatially overlapping activation during tool perception and action does not establish shared neural representations (Dinstein et al., 2008; Martin, 2007). We recently found that hand-selective, rather than tool-selective, visual regions represented action-relevant grasp information during interactions with 3D tools (Knights et al., 2021), suggesting sensitivity to learned object-hand interactions. Yet tool-use actions extend beyond grasp affordances to learned characteristic actions such as rotating a screwdriver or squeezing tongs. Whether neural responses during tool perception contain information about these actions, and whether this generalises to action production, remain open questions.

To address these questions, we conducted an fMRI study (Fig. 1) in which participants viewed familiar tools associated with different use actions (rotate or squeeze) and, separately, pantomimed tool-use actions to tool names. Tools differed in use action while being controlled for low-level visual properties and matched for grip type. Using multivoxel pattern analysis (MVPA), we first tested whether tool-use actions could be decoded during viewing and pantomime. Based on evidence for action-related representations within the tool-processing network, we predicted decoding in occipitotemporal and parietal regions implicated in tool perception and action (Mahon and Caramazza, 2009; Bracci et al., 2016; Striem-Amit et al., 2017). We then compared action with within-action tool-identity decoding (discrimination between tools matched for use action and grip type) to test whether these regions preferentially represented how tools are used rather than individual tool identity. Given evidence that hand-selective LOTC represents hand postures and action-relevant tool grasps (Bracci et al., 2018; Knights et al., 2021), we tested whether it contained relatively greater information about tool-use actions than individual tool identity. Finally, we tested whether action information generalised across viewing and pantomime. If tool perception recruits action representations shared with production of corresponding actions, action information should generalise across tasks.

**Figure 1.**
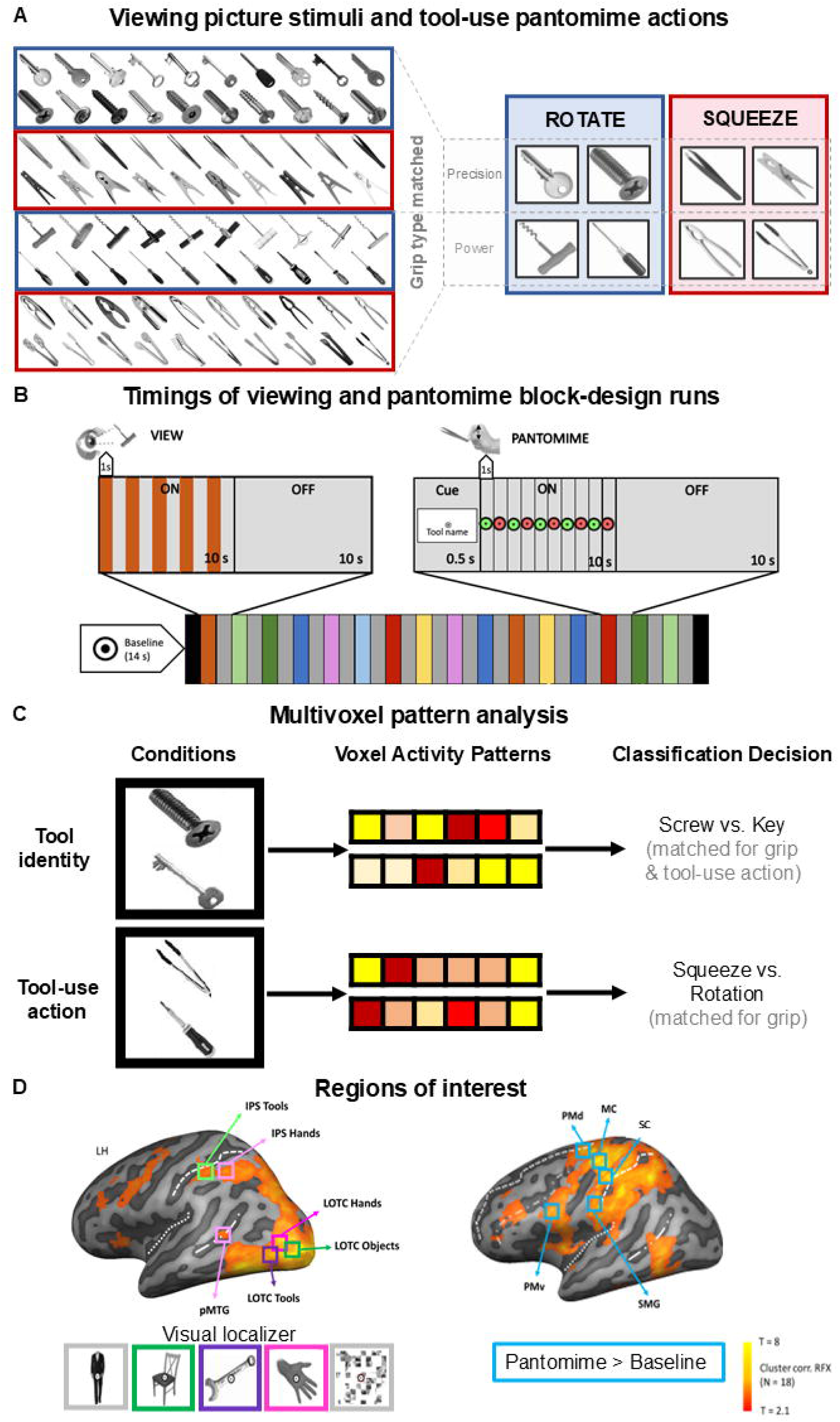
Experimental design and analysis. *A*, Tool stimuli presented during the viewing task (left) and corresponding tool-use actions performed during the pantomime task (right). Tools were associated with either rotate or squeeze actions and either precision or power grips, allowing decoding analyses to control for grip type. During pantomime, participants performed the corresponding tool-use action without the tool in hand, cued by the written tool name. ***B*,** Task sequence and timings. Viewing and pantomime tasks were completed in separate runs, counterbalanced across participants. Both tasks comprised 10-s ON and 10-s OFF blocks, with five stimulus presentations or pantomimed actions per ON block; pantomime blocks were additionally preceded by a 500-ms preparation cue containing the tool name. ***C*,** Decoding analysis. For each ROI and task separately, we performed tool-use action decoding (squeeze vs rotate) and within-action tool-identity decoding. Within-action tool-identity decoding discriminated individual tool pairs matched for both associated action and grip type (e.g., screw vs key: rotate/precision; tongs vs nutcracker: squeeze/power), ensuring that identity decoding could not be driven by action or grip differences. Tool-use action decoding controlled for grip type, with accuracies averaged across power- and precision-grip tool comparisons. ***D*,** ROI definition. Visual ROIs were defined independently for each participant using category-selective contrasts from the separate visual localizer experiment. Additional dorsal, premotor, and sensorimotor ROIs were defined using an orthogonal contrast of all pantomimed actions versus baseline. Representative ROI locations (Table 1) are displayed on a group activation map projected onto the left-hemisphere cortical surface of the COLIN27 Talairach reference brain.

**Table 1.**
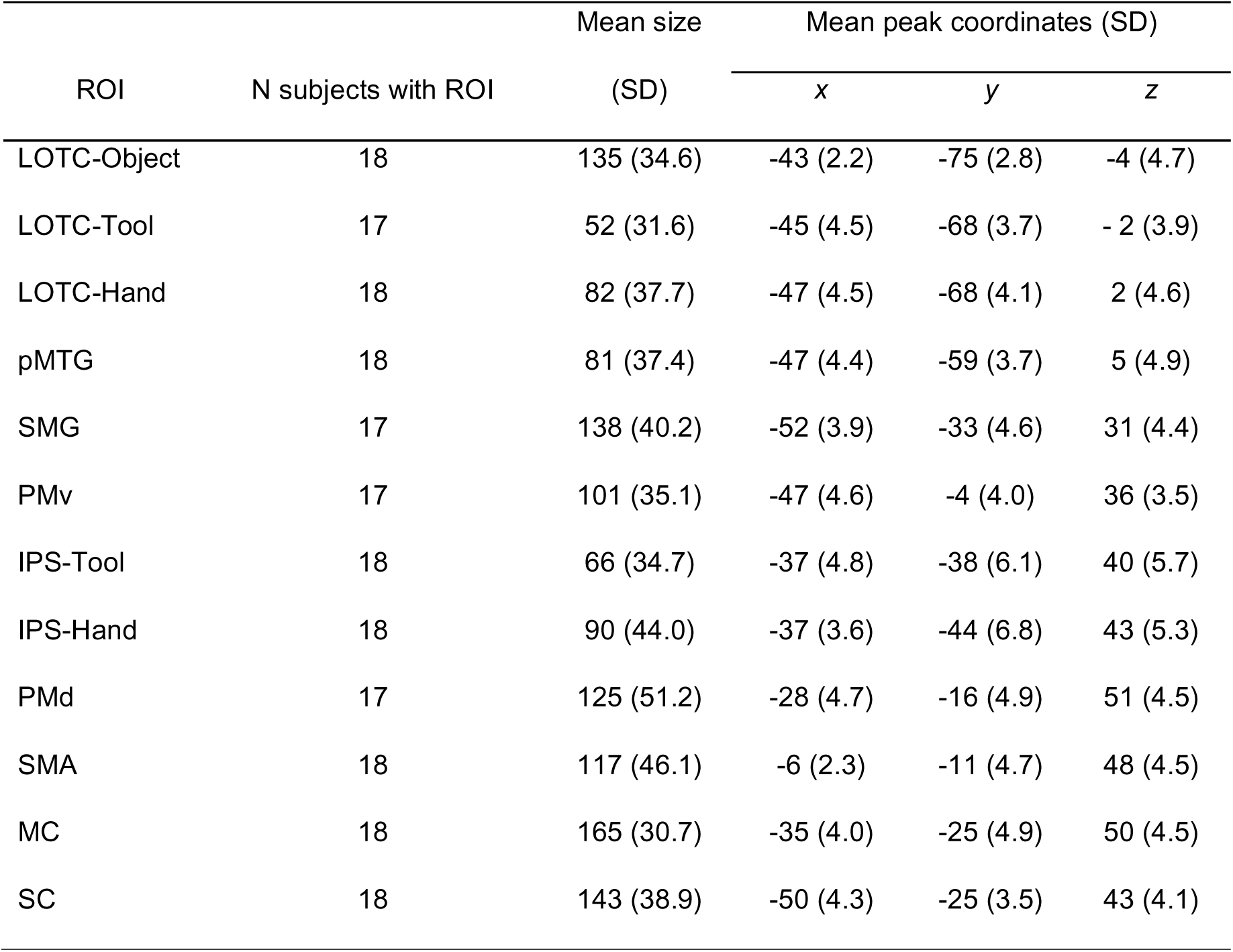
ROI descriptives. ROI subject counts with their mean sizes (voxels) and peak coordinates (Talairach).

## Materials and Methods

### Participants

Eighteen healthy participants aged 19-35 years [7 males; mean age = 24.7 (*SD* = 4.1)] recruited from the University of Maastricht (Netherlands) completed the main task and a visual localizer study in the same session. All participants had normal or corrected-to-normal vision and no history of neurological or psychiatric disorders, were right-handed (Oldfield, 1971), and provided informed consent in line with procedures approved by the School of Psychology Ethics Committee at the University of East Anglia. Our sample size was based on similar motor studies using MVPA (Ariani et al., 2015, 2018; Chen et al., 2016, 2018; Gallivan et al., 2013; Knights et al., 2021, 2022), though no power analysis was performed before data collection.

### Main experiment stimuli

The main experiment included viewing runs, in which participants viewed pictures of tools, and pantomime runs in which participants pantomimed tool-use actions (without tool in hand) in response to the names of the tools presented in the viewing runs. For the viewing task, ten exemplar images of each of eight tool identities were selected from the BOSS database (Brodeur et al., 2010, 2014), the Konklab database (Konkle and Oliva, 2011, 2012), and Google Images. To control for low-level visual differences between stimuli, all images were grayscale (800 x 800 pixels, app. 15°), presented on a white background and matched for luminance. Tool handles were all oriented rightwards at an angle of 45 degrees to control for any low-level visual effect of orientation. We chose to orient all tools to the right so that they would afford a right-handed action, given that all pantomime actions were performed with the right hand. To control for eye movements, participants were instructed to fixate on a bullseye (24 pixels, 0.5°) presented at the centre of the screen and superimposed on top of the centrally presented tool image. For the pantomime task, the English name of the tool was presented below the fixation bullseye in black on a white background (font size 32).

Our eight tool identities (tongs, screwdriver, tweezers, peg, key, screw, nutcracker, and corkscrew; see Fig. 1A) were selected from a larger dataset of 60 tools extracted from previous studies (Lagacé et al., 2013; Chen et al., 2016; Garcea and Mahon, 2012; McNair and Harris, 2012) that varied according to how the hand is moved to use the tool: half required a ‘rotate’ action (e.g., key and screwdriver), and the other half required a ‘squeeze’ action (e.g., peg and tongs).

Specifically, from the original 60 tools, we selected the ones that: 1) had similar grip types for using and grasping actions (Lagacé et al., 2013), ensuring that tools differed primarily in the tool-use action of interest (squeeze vs rotate), rather than grip type; 2) were associated with a single predominant tool-use action; 3) constituted a unique functional exemplar (e.g., spoon and wooden spoon are for the same function so only one was included); 4) afforded either a precision grip or power grip; and 5) when used did not involve interaction with the upper part of the body (e.g., lipstick, toothbrush, cup) or throwing actions to avoid excessive movements during pantomime actions in the scanner.

Using the twelve tools that fitted our criteria, we conducted a normative study at the University of East Anglia [N = 15, 1 male, mean age = 25.6 (*SD* = 6.5)] to match the final stimulus set in terms of familiarity and frequency of use. As such, our final eight tool identities were also consistently correctly named (*M* = 97%, *SD* = 5.55), highly familiar (median ratings > 6 on a 7-point Likert scale) and were rated as affording similar grasp types to both move and use the tool (median ratings > 5 on a 7-point Likert scale). To control for grip type differences, half of our final stimulus set afforded a precision grasp, and half afforded a power grasp. Analyses were run using grip-matched pairs (see Fig.1A).

### Main experiment fMRI set-up and paradigm

Participants lay supine in the scanner (using a standard coil configuration; see MRI acquisition) with their right upper arm restrained and supported by cushions so that pantomime movements were performed by flexion at the elbow thus reducing the likelihood of motion artifacts (Culham, 2006). Participants were instructed to maintain fixation on a central red bullseye throughout the task. To verify that the correct tool-use action was pantomimed we filmed their actions using a video camera (Panasonic HD HVC-210). The experiment was controlled using custom software written in MATLAB (MathWorks) using the Psychophysics Toolbox (Brainard, 1997). We used a powerful block-design fMRI paradigm (10s ON, 10s OFF; as in Knights et al., 2021), which maximized the contrast-to-noise ratio to generate a reliable estimate of the average response pattern (Mur et al., 2009) and improved detection of blood oxygenation level-dependent (BOLD) signal changes without significant interference from artifacts during overt movement (Birn et al., 2004). Viewing and pantomime tasks were completed in separate runs, counterbalanced across participants so that the same task was never completed in two consecutive runs. The action periods and OFF-blocks were matched across viewing and pantomime tasks (i.e., 10s ON, 10s OFF with 5 repetitions; Fig. 1B), with pantomime blocks additionally preceded by a 500-ms preparation cue.

In the viewing task, during the ON-block (10s), 5 image exemplars of a tool identity (e.g., 5 different pegs) were each presented for 1s with a blank inter-stimulus interval (ISI) of 1s. The ON-block was followed by a fixation OFF-block of 10s. The images were projected onto a screen (1920 x 1200 pixels) and viewed through a mirror mounted on the head coil (distance of mirror to screen = 60cm). Each run comprised 16 stimulus ON-blocks, with each tool identity being presented twice per run. Each exemplar was only presented once per block (thus 5 exemplars in one block, and the remaining 5 exemplars in another block). Each run started and ended with a 14s baseline block. Participants performed a 1-back repetition task by pressing their right index finger on the response box (2.5 x 13 x 6.4cm, fORP 932 response box system) whenever two successive blocks contained the same tool identity. Each viewing run lasted 348 seconds and participants completed on average 4.5 runs (minimum, four runs; maximum, five runs) for a total of 8-10 blocks per tool identity.

In the pantomime task, a block began with a preparation period (500ms) during which the tool name (e.g., tweezers) was presented cueing the tool-use action required (e.g., squeeze). Participants were instructed to lift their fist from the chest to prepare for the upcoming action. After this, the tool name disappeared, and fixation turned green 5 times (1s on, 1s off) to cue the participant to pantomime the tool-use action. At the end of each block, the fixation turned red to cue the participant to return their fist to their chest. Thus, in each ON-block (10s), five tool-use actions were pantomimed for 1s with a blank ISI of 1s. As in the viewing runs, each run comprised 16 stimulus ON-blocks, with each tool identity presented twice. A pantomime run lasted 356 seconds, slightly longer than the viewing run (given the extra 500ms required to present the tool names) and participants completed on average 4.66 runs (minimum, four runs; maximum, five runs) for a total of 8-10 blocks per tool identity.

Each main experiment session lasted ∼1.5 hours [including set-up and anatomical scan]. Immediately before the fMRI experiment, participants were familiarized with the stimuli and tasks (procedure adapted from Lausberg et al., 2015). Firstly, they were asked to name each tool image and demonstrate how they would pantomime their use. Then, the experimenter demonstrated the required tool-use action and participants repeated the pantomime in a comfortable and natural manner. Participants were then presented with the tool names and were required to pantomime the expected tool-use action when the word appeared on a screen until they reached 100% correct performance. Finally, participants completed up to eight trials of the pantomime task to familiarize themselves with the task. Three pantomime runs and two viewing runs were excluded from further analysis due to technical issues and excessive motion (i.e., translation and rotation exceeded 1.5 mm and 1.5°).

### Visual localizer

Following the main experiment, participants immediately completed a bodies, objects (chairs), tools, and hands visual localizer (Knights et al., 2021). Two sets of exemplar images were selected from previous stimuli databases (Bracci et al., 2012; Bracci and Op de Beeck, 2016; Bracci and Peelen, 2013) that were chosen to match, as much as possible, the characteristics within the tool (i.e., identity and orientation), body (i.e., gender, body position, and amount of skin shown), hand (i.e., position and orientation), and chair (i.e., materials, type, and style) categories. Different tool exemplars were used from the main fMRI experiment to avoid circularity. Using a mirror attached to the head coil, participants viewed separate blocks (14 s) of 14 grayscale 2D pictures from a given category (800□×□800 pixels; 0.5 s). Blank intervals separated individual stimuli (0.5 s), and scrambled image blocks separated cycles of the four randomized category blocks. Throughout the runs, participants fixated on a superimposed bullseye in the centre of each image and, to encourage attention, performed a one-back repetition detection task where they made a right-handed button press whenever two successive photographs were identical. A single fMRI run included 24 category blocks (six repetitions per condition) with blank fixation baseline periods (14 s) at the beginning and the end of the experiment. Each localizer scan lasted 448 s, and each participant completed at least 3 runs (*M* = 3.05, *SD* = 0.23) for a total of 18-24 repetitions per condition. The entire localizer lasted ∼30 min.

### MRI acquisition

BOLD fMRI measurements were acquired using a whole-body 3T scanner (MAGNETOM Prisma Fit, Siemens) with a 64-channel head coil at the Scannexus imaging center (Maastricht, The Netherlands). Functional images were acquired using a T2*-weighted echoplanar imaging sequence [35 horizontal slices, repetition time (TR) = 2000ms, echo time (TE) = 30ms, flip angle (FA) = 77°, field of view (FOV) = 216mm, matrix size (MS) = 72 x 72, voxel resolution = 3mm^3^]. High resolution T1-weighted anatomical images were also collected in the same session as functional scans (192 horizontal slices, TR = 2250ms, TE = 2.21ms, FA = 9°, FOV = 256mm, MS = 256 x 256, voxel size = 1mm^3^).

### Data pre-processing

Pre-processing and ROI definitions were performed using BrainVoyager QX (version 2.8; Brain Innovation). The BrainVoyager 3D motion correction (sinc interpolation) aligned each functional volume within a run to the functional volume acquired closest in time to the anatomical scan (Rossit et al., 2013). Slice scan time correction (ascending and interleaved) and high-pass temporal filtering (two cycles per run) were also performed. Functional data were superimposed onto the anatomical brain images that were previously aligned to the AC-PC plane and transformed into stereotaxic space (Talairach and Tournoux, 1988). Excessive motion was screened by examining the time-course movies and motion plots created with the motion-correction algorithms for each run. No spatial smoothing was applied.

To estimate activity in the localizer experiment, a predictor was used per image condition [i.e., bodies, objects (chairs), tools, hands, and scrambled] in a single-subject general linear model. Predictors were created from boxcar functions that were convolved with a standard 2*y* model of the hemodynamic response function (Boynton et al., 1996) and aligned to the onset of the stimulus with durations matching block length. The baseline epochs were excluded from the model, and therefore, all regression coefficients were defined relative to this baseline activity. This process was repeated for the main experiment, using 16 separate predictors for each block of stimulation independently per run and 6 motion regressors (confound predictors). These estimates (β weights) from the main experiment were used as the input to the pattern classifier.

### Regions of interest selection

Twelve regions of interest (ROIs) were defined at the individual participant level by drawing a cube (15mm^3^) centred on the peak of activity from previously reported volumetric contrasts (see below; Fig. 1D, Table 1) set at a threshold of *p* < 0.005 or, if no activity was initially identified, of p < 0.01 (Knights et al., 2021). In cases where no activity was observed at the liberal threshold, the ROI was omitted for that participant (Table 1). Given the predominantly left-lateralized nature of tool processing (Lewis, 2006), all individual participant ROIs were defined in the left hemisphere (Bracci et al., 2012; Bracci and Op de Beeck, 2016; Bracci and Peelen, 2013; Knights et al., 2021; Peelen et al., 2013). Data from the independent visual localizer was used to define 6 visual regions of interest (ROIs) in the left hemisphere: lateral occipitotemporal cortex (LOTC)-object, LOTC-tool, LOTC-hand, posterior middle temporal gyrus (pMTG), intraparietal sulcus (IPS)-hands, and IPS-tool. Two tool-selective ROIs were identified by contrasting tool pictures versus other objects or scrambled pictures [IPS-tool; LOTC-tool]. Moreover, two hand-selective ROIs were identified in LOTC (LOTC-hand) and IPS (IPS-hand) by contrasting activation for hand pictures versus pictures of other body parts (Bracci et al., 2012, 2018; Peelen et al., 2013; Bracci and Op de Beeck, 2016; Palser and Cavina-Pratesi, 2018), and a hand-selective region was identified in pMTG by contrasting hand pictures with pictures of other objects (we were unable to find reliable activation in pMTG using the traditional tools > chairs contrast; Knights et al., 2021). Additionally, we defined an object-selective ROI [LOTC-object selective (LOTC-object); chairs > scrambled; Bracci and Op de Beeck, 2016; Hutchison et al., 2014]. Finally, an orthogonal contrast was applied to pantomime runs (all actions > baseline; Gallivan et al., 2011) to define tool processing ROIs in the dorsal-dorsal and ventro-dorsal streams [i.e., supramarginal gyrus (SMG); ventral premotor cortex (PMv); dorsal premotor cortex (PMd)] and sensorimotor cortices [supplementary motor area (SMA); motor cortex (MC); somatosensory cortex (SC)]. This contrast was orthogonal to the distinctions subsequently used for decoding and therefore did not select voxels based on differences between action or identity conditions (Kriegeskorte et al., 2009). The ROI locations were verified by a senior author (S.R.) with respect to the following anatomical guidelines and contrasts:

*LOTC-object (*chairs > scrambled; Bracci and Op de Beeck, 2016; Knights et al., 2021) is defined by selecting the peak of activation near the lateral occipital sulcus (LOS; Hutchison et al., 2014; Bracci and Op de Beeck, 2016; Malach et al., 1995; Grill-Spector et al., 1999, 2001).

*LOTC-hand* [(hands > chairs) and (hands > bodies); Bracci and Op de Beeck, 2016] is defined by selecting the peak of activation near the LOS. This peak was often anterior to LOTC-body (bodies > chairs; Bracci et al., 2010; Bracci and Op de Beeck, 2016). LOTC-body was not included in the analysis.

*LOTC-tool* (tools > chairs; Bracci et al., 2012; Hutchison et al., 2014) is defined by selecting peak of activation near the LOS which closely overlapped with LOTC-hand (Bracci et al., 2012).

*pMTG* (hands > chairs) is defined by selecting the peak of activation on the pMTG, more lateral, ventral and anterior to extrastriate body area (Hutchison et al., 2014). We selected the peak anterior to the anterior occipital sulcus (AOS), as the MTG is in the temporal lobe and the AOS separates the temporal lobe from the occipital lobe (Damasio, 1995).

*SMG* (pantomimes > baseline) is defined by selecting the peak of activation along the SMG, lateral to the anterior portion of the IPS (Gallivan et al., 2013).

*PMv* (pantomimes > baseline) is defined by selecting the voxels inferior and posterior to the junction between the inferior frontal sulcus and pre-central sulcus (Gallivan et al., 2013).

*IPS-tool* (tools > scrambled; Bracci and Op de Beeck, 2016) is defined by selecting the peak of activation close to the junction between the anterior intraparietal sulcus (aIPS) and post-central sulcus.

*IPS-hand* (hands > chairs; Bracci and Op de Beeck, 2016) is defined by selecting the peak of activation along the IPS (Bracci and Op de Beeck, 2016).

*PMd* (pantomimes > baseline) is defined by selecting the peak of activation at the junction of the precentral sulcus and the superior frontal sulcus (Gallivan et al., 2013).

*SMA* (pantomimes > baseline; Fabbri et al., 2014) is defined by selecting the peak of activation on the medial wall of the posterior frontal gyrus, anterior to the medial end of the central sulcus and posterior to the vertical projection of the anterior commissure plane (Ariani et al., 2015).

*MC* (pantomimes > baseline) is defined by selecting the peak of activation around the “hand knob” area in the anterior bank of the central sulcus (Gallivan et al., 2013).

*SC* (pantomimes > baseline; Fabbri et al., 2014) is defined by selecting the peak of activation medial and anterior to the aIPS, including the post-central gyrus and post-central sulcus (Gallivan et al., 2013).

### ROI within-task decoding

MVPA was performed separately for viewing and pantomime runs. Independent linear pattern classifiers [linear support vector machine (SVM)] were trained to learn the mapping between a set of brain-activity patterns (β values computed from single blocks of activity) from the ROIs and the individual tool identities or tool-use actions either depicted by the pictures (in the viewing task) or pantomimed tool-use actions (in the pantomime task). Specifically, for each ROI and task separately, we performed within-action tool-identity decoding (i.e., pairwise discrimination between individual tools matched for tool-use action and grip type) and tool-use action decoding (squeeze vs rotate), while controlling for grip-type differences between tool pairs (Fig.1C). Following this, the decoding accuracies were averaged across the power and precision tools, ensuring that tool-use action decoding was not confounded by grip-type differences. For within-action tool-identity decoding, the classifier was trained and tested to distinguish between tool pairs that differed in individual tool identity but were matched for tool-use action and grip type the tool was associated with (i.e., separate decoding analyses were run for tool pairs associated with different actions; see Fig. 1C). For example, the classifier was trained to distinguish between screw and key (both requiring a rotate action, and precision grip), and between tongs and nutcracker (both requiring a squeeze action, and power grip). Thus, tool-identity decoding was performed within action and grip categories and could not be driven by differences in tool-use action or grip type. Given the two-class classification procedure, chance-level decoding accuracy was 50%.

To test the performance of our classifiers, decoding accuracy was assessed using an *n*-fold leave-one-run-out cross-validation procedure; thus, our models were built from *n* - 1 runs and were tested on the independent *n*th run (repeated for the n different possible partitions of runs in this scheme; Duda et al., 2001; Smith and Muckli, 2010; Smith and Goodale, 2015; Gallivan et al., 2016; Knights et al., 2021; Bailey at al., 2023) before averaging across *n* iterations to produce a representative decoding accuracy measure per participant and per ROI. Beta estimates for each voxel in the training data were normalized within a range of −1 to +1 before input to the SVM (Chang and Lin, 2011) and the test data normalized to the same scale, and the linear SVM algorithm was implemented using the default parameters provided in the LibSVM toolbox (C□=□1). Pattern classification was performed with a combination of in-house scripts (Smith and Muckli, 2010; Smith and Goodale, 2015; Bailey et al., 2023) using MATLAB with the Neuroelf toolbox (version 0.9c; http://neuroelf.net) and a linear SVM classifier (libSVM 2.12 toolbox; https://www.csie.ntu.edu.tw/∼cjlin/libsvm/).

For each ROI and task, one-tailed one-sample *t* tests were used to test for above-chance decoding for tool-use action and tool identity classifications (missing data were replaced with task and condition mean). Additionally, to compare sensitivity to tool-use action and individual tool identity, we contrasted action and identity decoding within each task and ROI by running a 2 x 2 x 12 repeated-measures ANOVA with property decoded (tool identity, tool-use action), task (view, pantomime), and ROI (LOTC-object, LOTC-hand, LOTC-tool, pMTG, IPS-hand, IPS-tool, SMG, SC, SMA, PMv, PMd, MC) as within-subject factors. For all analyses, we corrected for multiple comparisons with false discovery rate (FDR) correction of *q* ≦ 0.05 across the number of tests (Benjamini and Hochberg, 1995; Benjamini and Yekutieli, 2001). To determine whether decoding effects could be explained by differences in mean response amplitude, we conducted a complementary univariate analysis using β-weight difference scores matched to the comparisons used for MVPA (see Supplementary Materials).

### ROI cross-task decoding

To test whether action-specific information generalised across tool viewing and pantomime, we conducted cross-task decoding by training the classifier on activity patterns from the viewing task and testing it on activity patterns from the pantomime task, and vice versa, using the leave-one-run-out procedure described above. Performance was averaged across the two possible training and test directions. If participants had a greater number of runs in one task than another, we reduced the number of runs in the task with more to match via run number (if P1 had 5 runs in task 1 and 4 in task 2, we removed the 5th run from the task 1 data to ensure parity). Note that this analysis used the same procedures as the within-task decoding to control for grip type in tool-use action decoding and for grip type and tool-use action in tool-identity decoding. As in our within-task MVPA, we conducted a series of one-tailed one-sample t-tests against chance in each of our ROIs. All tests were FDR corrected to control for multiple comparisons (*q* ≦ 0.05).

### Whole-brain searchlight decoding

To identify regions across the whole brain containing information about tool-use action and individual tool identity during viewing and pantomime, we performed whole-brain searchlight MVPA independently, per participant, using separate linear pattern classifiers for the viewing and pantomime tasks (as in the within-task decoding above). A cube mask (5 x 5 x 5 voxel length, 125 voxels) was shifted through the entire brain volume, applying the classification procedure at each centre voxel (Smith and Goodale, 2015; Knights et al., 2022) to measure the accuracy that a given cluster of activity patterns could be used to discriminate between different tool-use actions or individual tool identities within either the viewing or pantomime tasks. We also conducted further cross-decoding analyses (as above). To test performance of the classifiers, for within-task decoding we again used leave one-run out cross-validation (as above) but for the cross-task decoding, for computational efficiency, we used all available data from one task to train the classifier and all data from the other task to test the classifier. Regardless, each analysis produced a decoding accuracy value per voxel for each analysis. Searchlight analysis space was restricted to a common group mask within Talairach space, defined by voxels with a mean BOLD signal□>□100 for every participant’s fMRI runs to ensure that all voxels included in searchlight MVPA contained suitable activation. As in the ROI decoding analyses, beta estimates for each voxel were normalized (separately for training and test data) within a range of□−□1 to 1 before input to the SVM, and the linear SVM algorithm was implemented using the default parameters provided in the LibSVM toolbox (C□=□1). Pattern classification was performed with a combination of in-house scripts (Knights et al., 2022; Smith and Goodale, 2015; Bailey et al., 2023) implemented in Matlab using the SearchMight toolbox (Pereira and Botvinick, 2011).

Searchlight MVPA produced unsmoothed participant-level decoding-accuracy maps. To identify brain regions containing above-chance information about individual tool identity or tool-use action, we used a one-sample t-test (across participants) against the chance level performance (50%; see Smith and Goodale, 2015). The BrainVoyager cluster-level statistical threshold estimator (Goebel et al., 2006; Forman et al., 1995) was used for cluster correction (voxelwise thresholds were set to *p□*=□0.01 and then the cluster-wise thresholds were set to *p□*<□0.05 using a Monte Carlo simulation of 1000 iterations), before projecting results on to a standard surface (Xia et al., 2013).

### Data availability

Stimuli, code for running the experiment and for MVPA analyses, and ROI data are accessible from Open Science Framework at https://osf.io/md53t/.

## Results

### ROI decoding

We first tested which ROIs showed above-chance (>50%; FDR-corrected) tool-identity decoding (within action) during picture viewing and tool-use pantomime. As shown in Figure 2, LOTC-hand and SMG were the only ROIs in which tool identity was decoded significantly above chance in both the viewing and pantomime tasks [mean ± SD (LOTC-hand accuracies: viewing = 58.16 ± 7.49%, *t*(17) = 4.62, *p* < .001, *d* = 1.09; pantomime = 52.71 ± 5.88%, *t*(17) = 1.96, *p* = .034, *d* = .461); SMG accuracies: viewing = 53.47 ± 7.23%, *t*(17) = 2.04, *p* = .029, *d* = .481; pantomime = 54.49 ± 5.63%, *t*(17) = 3.38, *p* = .002, *d* = .797)]. Beyond LOTC-hand and SMG, which showed above-chance tool identity decoding in both tasks, task-specific differences were apparent across ventral and dorsal ROIs. During viewing, additional above-chance decoding was observed in ventral ROIs including LOTC-object, LOTC-tool, and pMTG, whereas during pantomime, additional above-chance decoding was observed in dorsal and sensorimotor ROIs including IPS-tool, IPS-hand, PMd, SC, and MC [viewing LOTC-object accuracy = 62.57 ± 8.83%, *t*(17) = 6.04, *p* < .001, *d* = 1.42; viewing LOTC-tool accuracy = 59.72 ± 8.10%, *t*(17) = 5.10, *p* < .001, *d* = 1.20; viewing pMTG accuracy = 55.42 ± 7.09%, *t*(17) = 3.24, *p* = .002, *d* = 0.764); pantomime IPS-tool accuracy = 54.60 ± 6.97%, *t*(17) = 2.80, *p* = .006, *d* = 0.660; pantomime IPS-hand accuracy = 55.10 ± 6.37%, *t*(17) = 3.40, *p* = .002, *d* = .801; pantomime MC accuracy = 60.35 ± 9.11%, *t*(17) = 4.82, *p* < .001, *d* = 1.14; pantomime SC accuracy = 62.34 ± 9.53%, *t*(17) = 5.50, *p* < .001, *d* = 1.30, pantomime PMd accuracy = 57.02 ± 9.25%, *t*(17) = 3.22, *p* = .003, *d* = .759].

**Figure 2.**
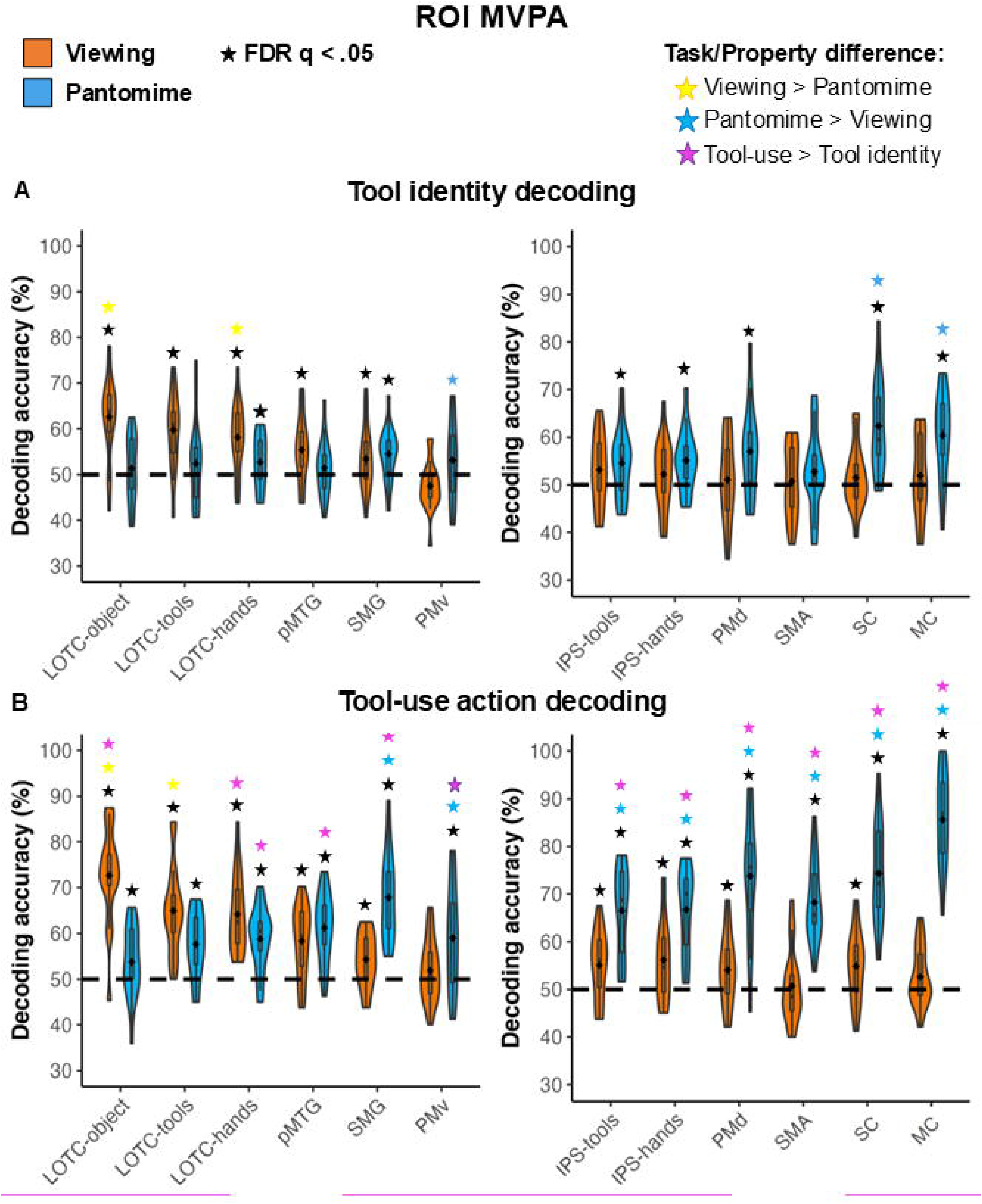
Within-task decoding in left-hemisphere ROIs. Violin plots show decoding accuracy for ***A***, within-action tool identity and ***B***, tool-use action during viewing and pantomime in ventral/ventro-dorsal (left) and dorsal-dorsal/sensorimotor (right) ROIs. Tool-identity decoding discriminated tools matched for associated action and grip type; tool-use action decoding discriminated squeeze from rotate actions while controlling for grip type. Box-plot centre lines indicate mean decoding accuracy; edges and whiskers indicate ±1 SD and ±2 SEM, respectively. The dashed line indicates chance-level decoding (50%). Black stars indicate significant above-chance within-task decoding; coloured stars indicate significant post-hoc comparisons following the property decoded × task × ROI ANOVA, comparing decoding between tasks or between tool-use action and tool identity within each ROI. All statistical comparisons were FDR-corrected (q ≤ 0.05).

Next, we examined tool-use action decoding for both tasks (Table 2; Fig. 2). In line with our predictions, during viewing, tool-use action was decoded significantly above chance in hand- and tool-selective LOTC and IPS, as well as LOTC-object, pMTG, SMG, PMd, and even SC. During pantomime, tool-use action was decoded significantly above chance in all ROIs, even in ventral visual areas such as LOTC.

**Table 2.**
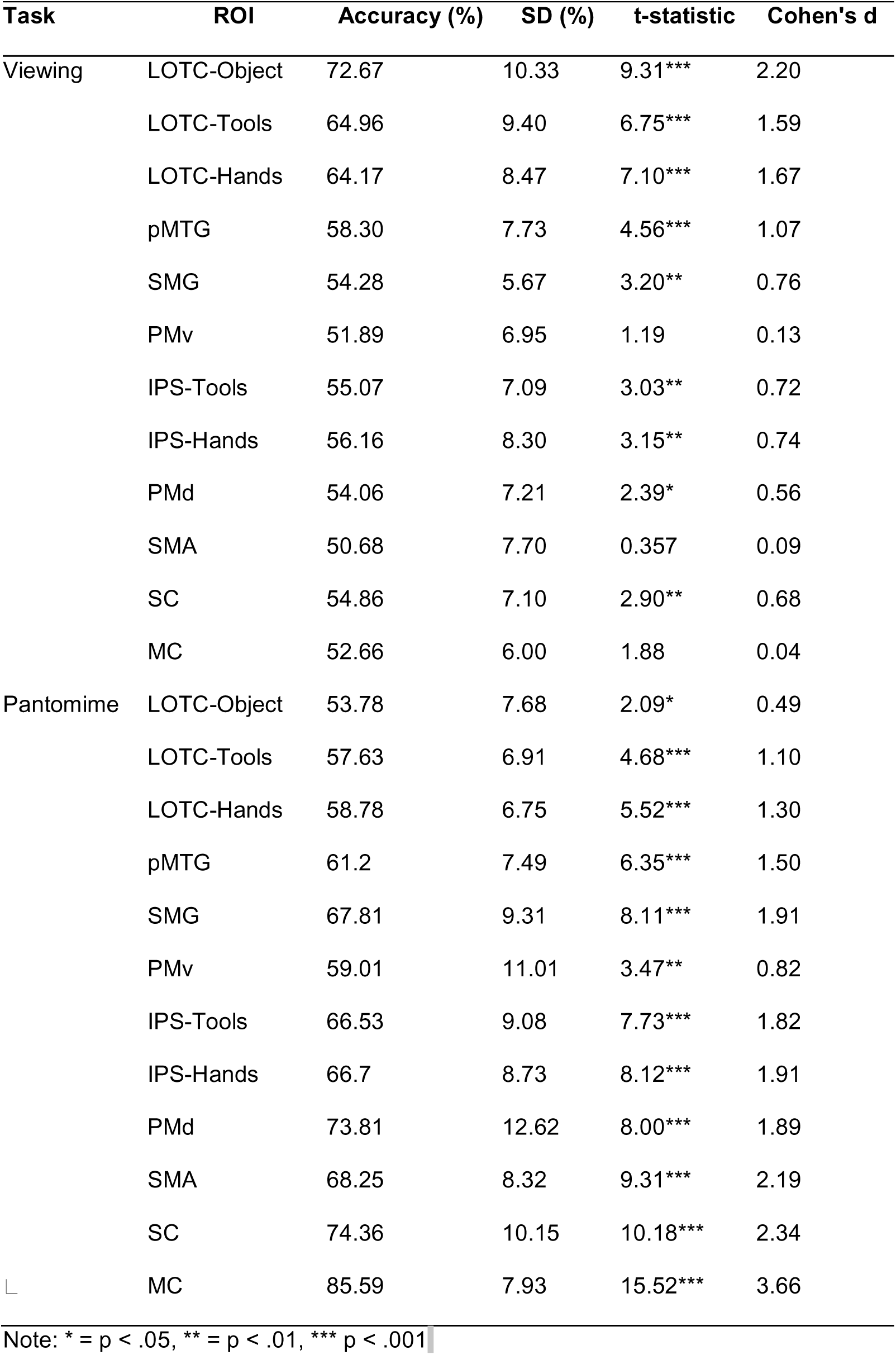
Tool-use decoding accuracies and effect sizes for all ROIs for viewing and pantomime tasks.

We next compared sensitivity to tool-use action with sensitivity to individual tool identity for each task and ROI and ran a 2 (property decoded: individual tool identity, tool-use action) × 2 (task: viewing, pantomime) × 12 (ROI) repeated-measures ANOVA. This revealed a significant three-way interaction [*F*(11, 187) = 6.293, *p* < .001, η_p_^2^ = .270]. Post-hoc tests (FDR corrected; see Table 3) showed that LOTC-hand was the only region in which tool-use action decoding exceeded individual tool identity decoding in both viewing and pantomime. Thus, LOTC-hand showed greater sensitivity to the action category associated with the tools than to the matched within-action tool-identity distinctions used here. Post-hoc comparisons also revealed task-dependent differences across ROIs, with generally higher decoding in ventral visual regions during viewing and higher decoding in dorsal and sensorimotor regions during pantomime (see Table 4).

**Table 3.** Regions showing a significant difference between tool-use action and tool identity decoding accuracy for viewing and pantomime tasks.

| Task | ROI | Comparison | Mean Difference (SE; %) | *p* |
| --- | --- | --- | --- | --- |
| Viewing | LOTc-object | Use Action > Identity | 10.10 (2.43) | .001 |
|  | LOTc-hand | Use Action > Identity | 6.01 (2.06) | .010 |
|  | PMv | Use Action > Identity | 4.38 (1.65) | .017 |
| Pantomime | LOTc-hand | Use Action > Identity | 6.08 (1.96) | .007 |
|  | pMTG | Use Action > Identity | 9.77 (1.91) | < .001 |
|  | SMG | Use Action > Identity | 13.33 (2.63) | < .001 |
|  | IPS-tool | Use Action > Identity | 11.93 (2.26) | < .001 |
|  | IPS-hand | Use Action > Identity | 11.60 (1.81) | < .001 |
|  | PMd | Use Action > Identity | 16.78 (3.57) | < .001 |
|  | SMA | Use Action > Identity | 15.54 (2.45) | < .001 |
|  | SC | Use Action > Identity | 12.01 (2.71) | < .001 |
|  | MC | Use Action > Identity | 25.24 (3.88) | < .001 |

**Table 4.** Regions in which decoding accuracy for tool identity and tool-use action significantly differed between viewing and pantomime tasks.

| Decoding Type | ROI | Comparison | Mean Difference (SE; %) | *p* |
| --- | --- | --- | --- | --- |
| Tool Identity | LOTc-object | Viewing > Pantomime | 11.22 (2.76) | < .001 |
|  | LOTc-hand | Viewing > Pantomime | 5.45 (1.85) | .009 |
|  | PMv | Pantomime > Viewing | 5.64 (2.02) | .012 |
|  | SC | Pantomime > Viewing | 10.85 (2.81) | .001 |
|  | MC | Pantomime > Viewing | 8.41 (2.74) | .007 |
| Tool-use action | LOTc-object | Viewing > Pantomime | 18.89 (2.37) | < .001 |
|  | LOTc-tool | Viewing > Pantomime | 7.34 (2.46) | .008 |
|  | SMG | Pantomime > Viewing | 13.53 (2.25) | < .001 |
|  | PMv | Pantomime > Viewing | 7.11 (2.69) | .017 |
|  | IPS-tool | Pantomime > Viewing | 11.46 (3.00) | .001 |
|  | IPS-hand | Pantomime > Viewing | 10.54 (2.83) | .002 |
|  | PMd | Pantomime > Viewing | 19.74 (3.16) | < .001 |
|  | SMA | Pantomime > Viewing | 17.56 (2.34) | < .001 |
|  | SC | Pantomime > Viewing | 19.50 (2.80) | < .001 |
|  | MC | Pantomime > Viewing | 32.93 (2.91) | < .001 |

Specifically, for tool identity decoding, LOTC-object and LOTC-hand showed significantly higher decoding during viewing than pantomime, whereas the opposite was true for PMv, SC and MC. Similarly, for tool-use action decoding, ventral regions LOTC-object and LOTC-tool showed higher decoding during viewing than pantomime, whereas SMG, PMv, IPS-tool, IPS-hand, PMd, SMA, SC and MC showed higher decoding during pantomime than viewing. Finally, beyond LOTC-hand, where tool-use action decoding exceeded tool-identity decoding in both tasks, task-specific effects were observed. Tool-use action decoding exceeded within-action tool-identity decoding only during pantomime in pMTG, SMG, IPS-tool, IPS-hand, PMd, SMA, SC and MC, and only during viewing in LOTC-object and PMv.

Finally, we tested whether the decoding effects could be explained by differences in mean response amplitude within each ROI (Fig. 3). Using β-weight difference scores matched to the comparisons used for MVPA (see Supplementary Materials), we observed a significant ROI × task interaction (*F*(4.86, 82.59) = 2.455, *p* = .041, η_p_^2^ = .126) and a significant ROI x property interaction (*F*(11, 187) = 2.168, *p* = .018, η_p_^2^ = .113). FDR corrected post-hoc comparisons revealed a larger difference in activation between the tool identities in IPS-hand compared to PMd (mean difference = .051, *SE* = .011, *p* < .001), and that in PMv, there was a greater activation difference between tool identity than tool-use action (mean difference = .049, *SE* = .010, *p* < .001). Thus, the pattern of univariate activation differences did not mirror the multivoxel decoding results, indicating that the decoding effects were not simply attributable to differences in mean response amplitude.

**Figure 3.**
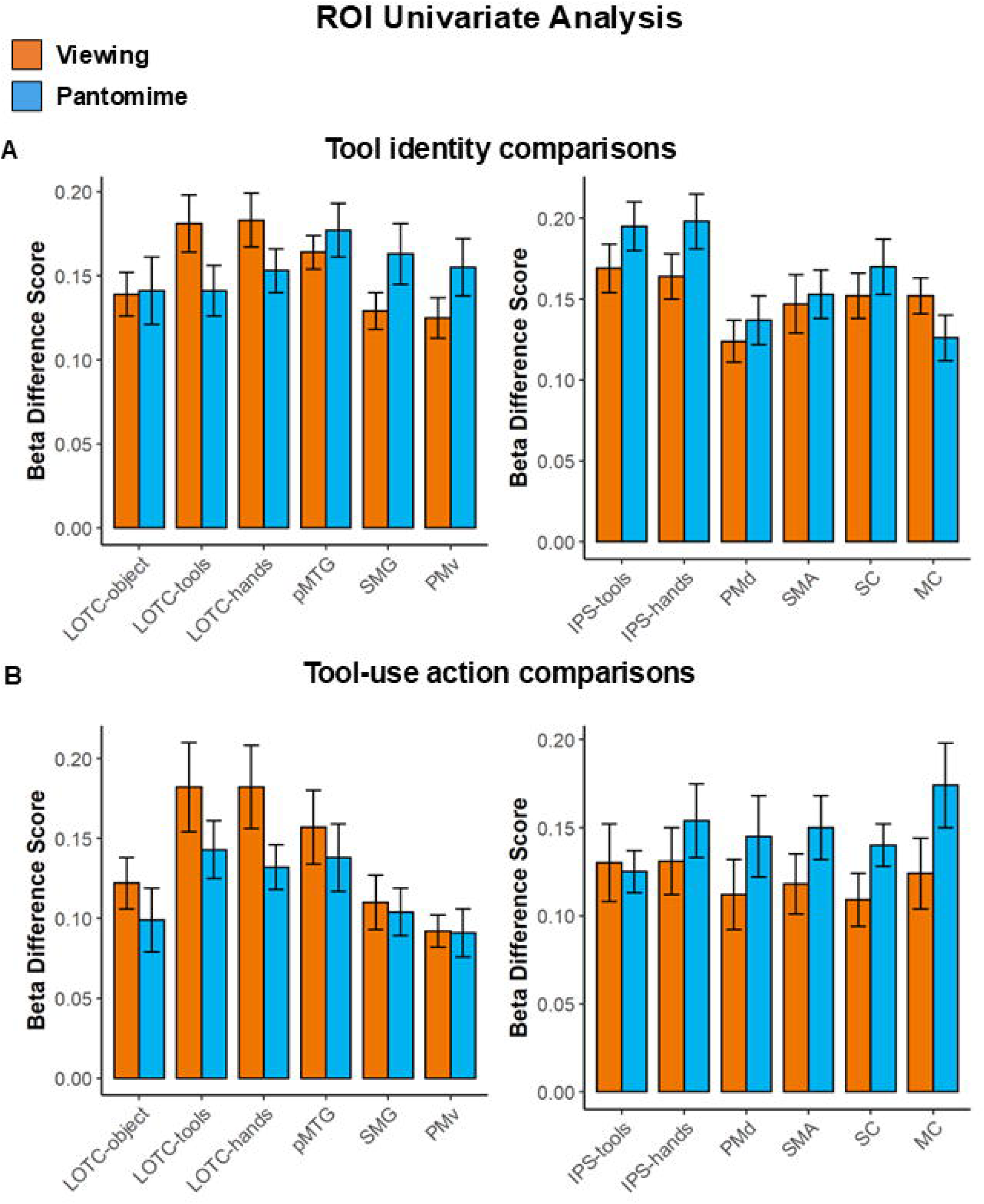
Univariate response differences corresponding to the MVPA comparisons. β-weight difference scores matched to the ***A***, tool-identity and ***B***, tool-use action comparisons used in the multivoxel decoding analyses. Error bars represent ±1 SEM.

### ROI cross-task decoding

Finally, we tested whether action-specific information generalised across tool viewing and pantomime by training classifiers on activity patterns from one task and testing them on the other. No ROI showed reliable above-chance cross-task decoding of tool-use action, providing no evidence that action-specific activity patterns were shared across viewing and pantomime.

### Whole-brain within-task searchlight

Results from the whole-brain searchlight analysis broadly converged with the ROI findings (Table 5; Fig. 4). Tool identity was decoded significantly above-chance during the viewing task in a large area of occipital cortex, including early visual cortex (EVC), extending bilaterally to ventral stream regions LOTC, fusiform gyrus and MTG. Significant decoding was also observed in dorsal regions bilaterally along IPS and SPOC. Posterior clusters of significant decoding were larger in the right than left hemisphere. Above-chance decoding of tool-use action in the viewing task was observed in a large area of the occipital cortex bilaterally, this extended to ventral regions LingG, LOTC, FusiG bilaterally, and left MTG. In dorsal regions, significant above-chance tool-use action decoding was observed in right IPS and left MC. We also observed above-chance decoding in bilateral SC.

**Figure 4.**
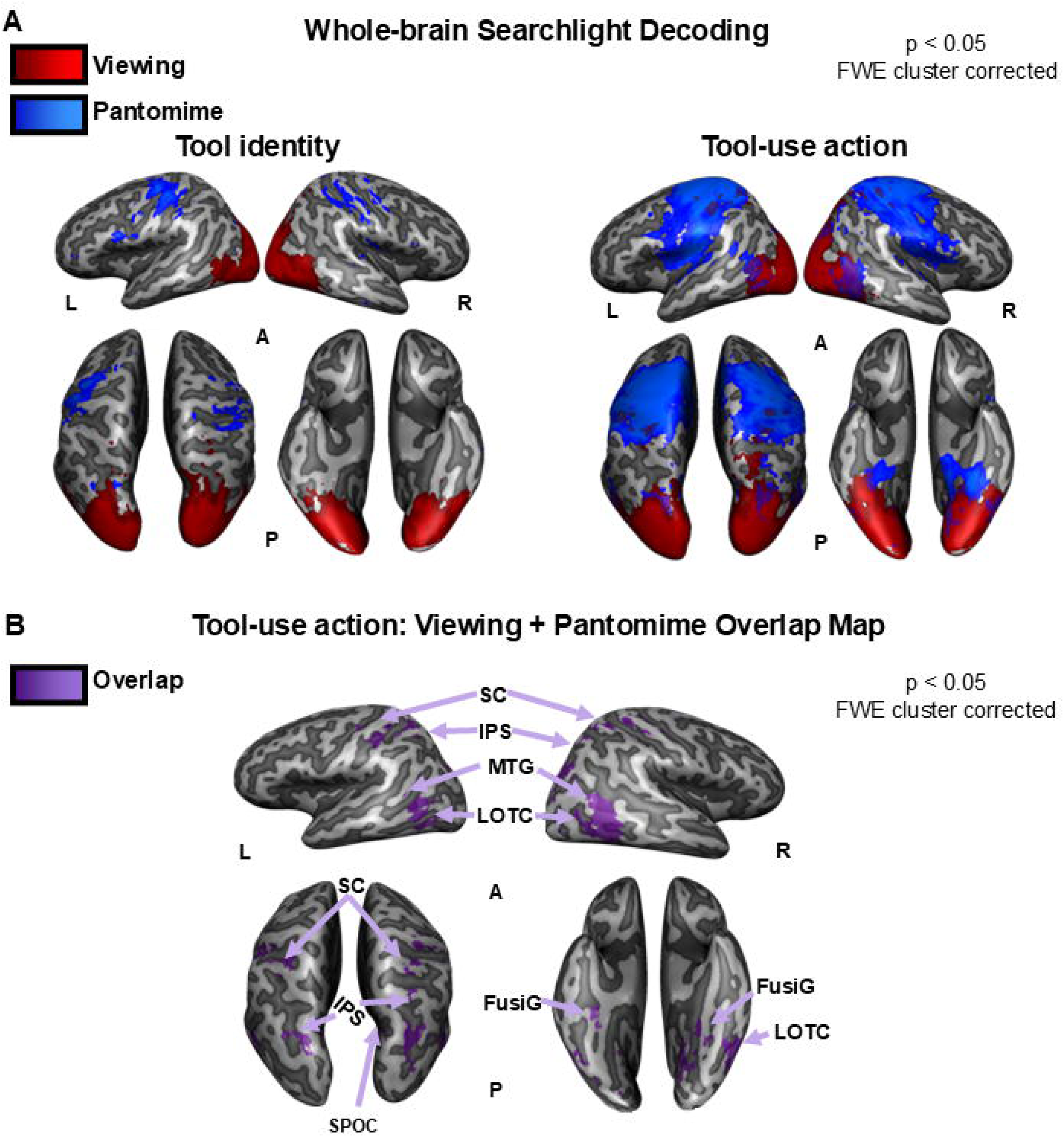
Whole-brain searchlight decoding. ***A***, Group maps showing significant above-chance within-task decoding of tool identity and tool-use action during viewing and pantomime. During viewing, decoding was predominantly distributed across posterior occipitotemporal and parietal regions, whereas pantomime particularly tool-use action decoding - produced more widespread frontoparietal and sensorimotor decoding. ***B***, Spatial overlap between regions showing significant within-task tool-use action decoding during viewing and pantomime, including occipitotemporal, parietal, motor, and somatosensory regions. Despite this anatomical overlap, no clusters showed significant cross-task decoding, providing no evidence that tool-use action information was represented in a shared multivoxel format across viewing and pantomime. Abbreviations: LOTC, lateral occipitotemporal cortex; FusiG, fusiform gyrus; MTG, middle temporal gyrus; SPOC, superior parieto-occipital cortex; IPS, intraparietal sulcus; SC, somatosensory cortex.

**Table 5.**
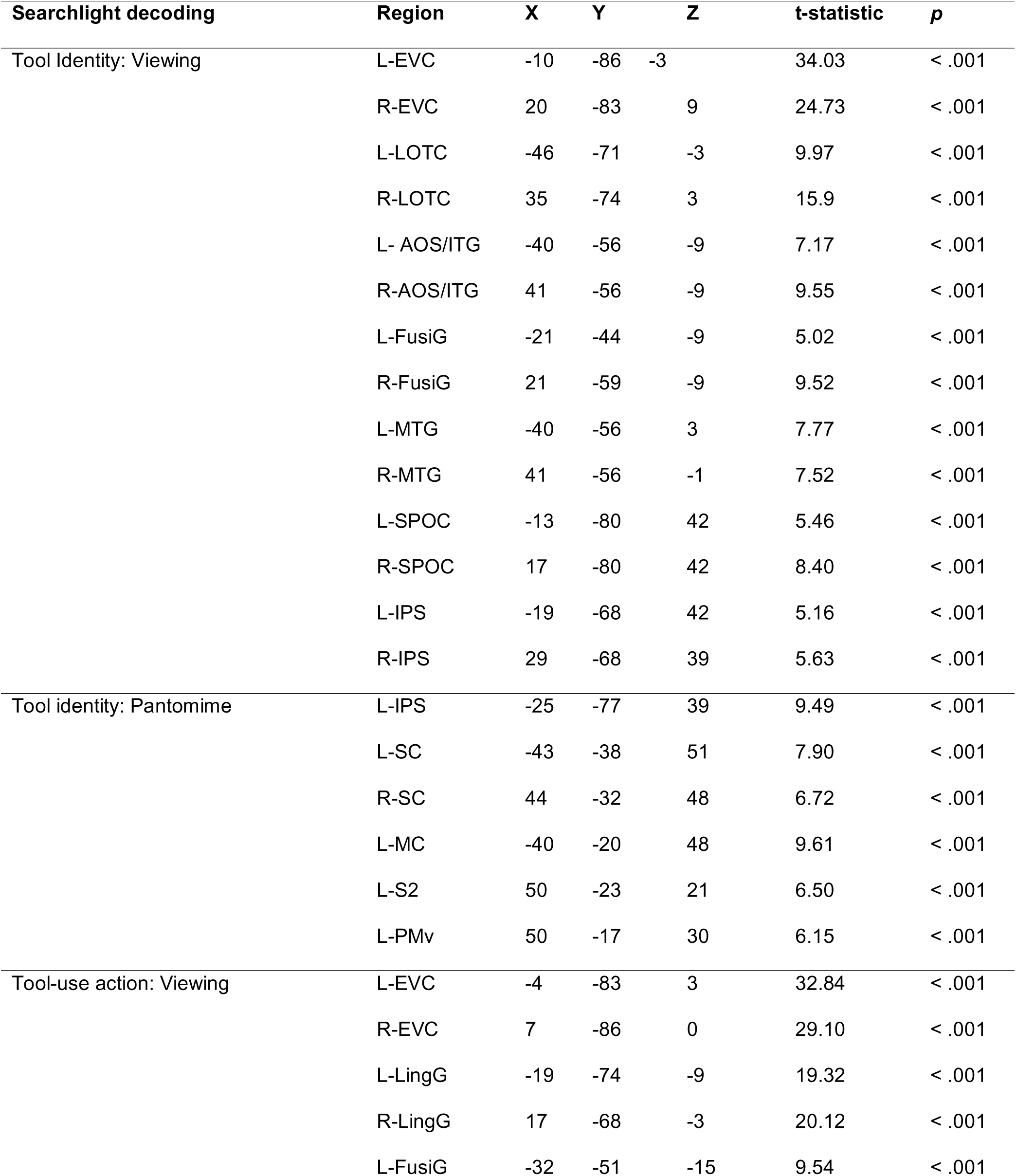
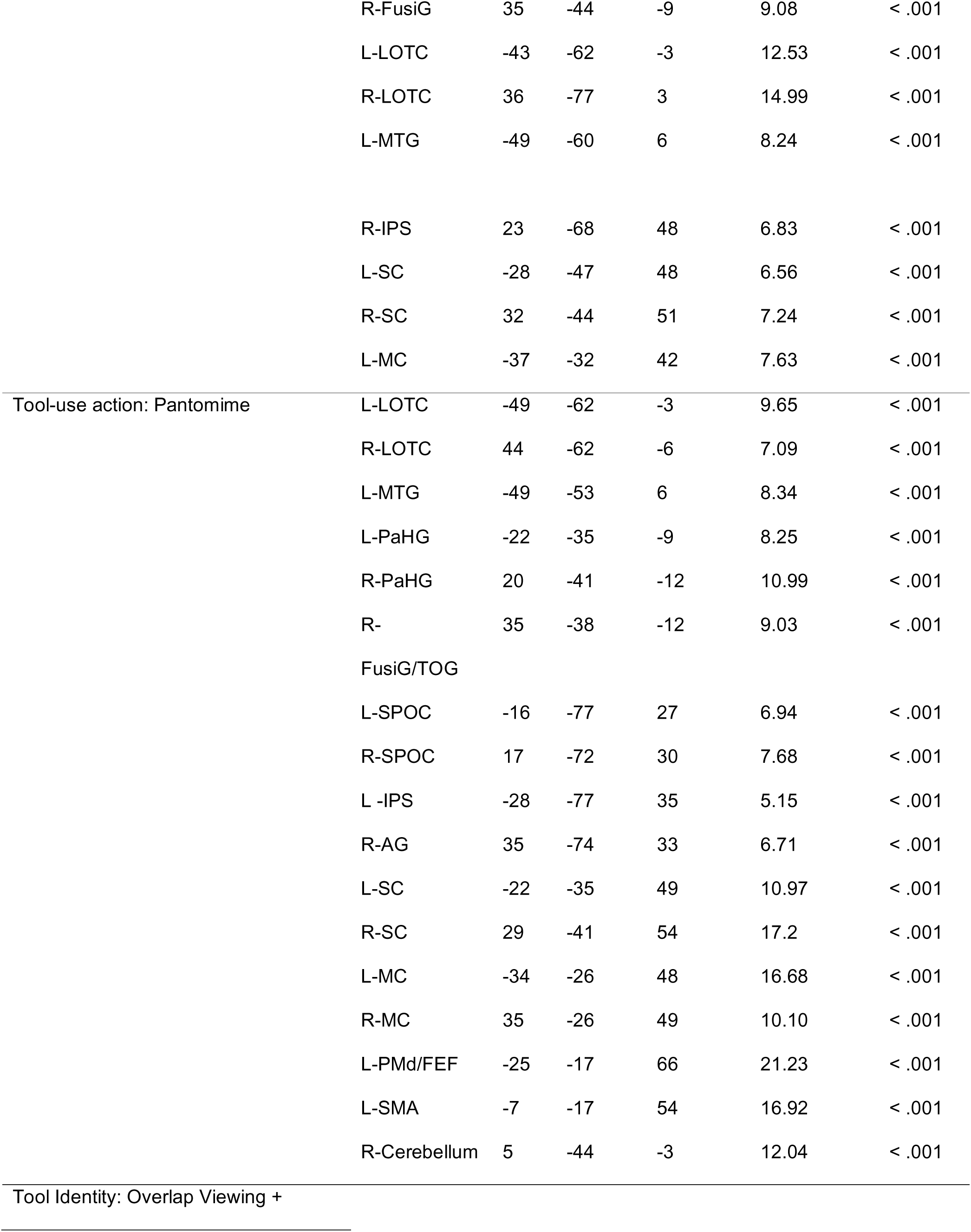
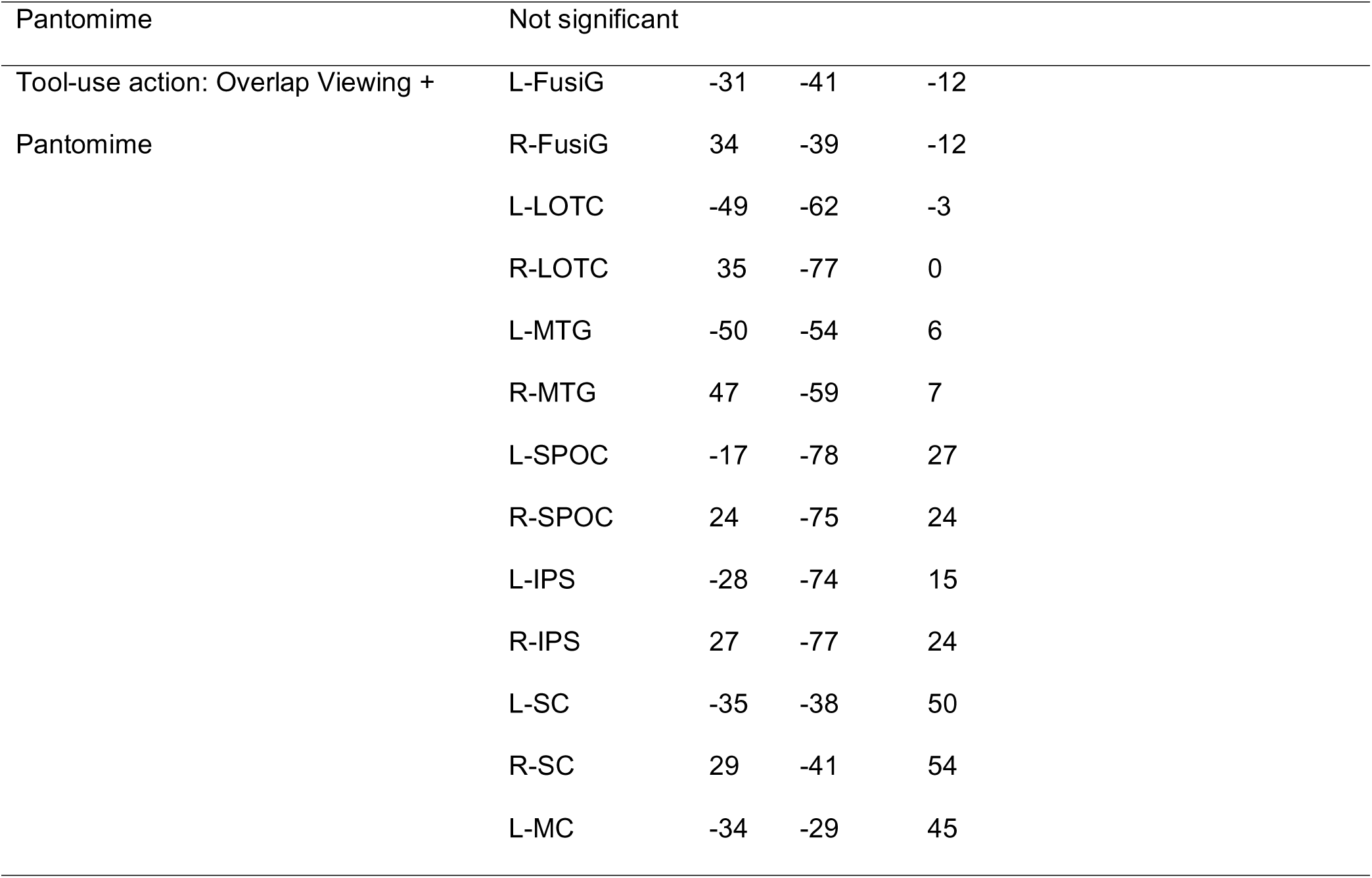
Searchlight peak coordinates (Talairach) and statistics for decoding tool identity and tool-use action categories for viewing and pantomiming tasks.

For the pantomime task, tool identity was successfully decoded in MC, SC and PMv. This was largely left-lateralised; however, a cluster was also found in right SC. Decoding of tool-use action in the pantomime task was also observed in bilateral motor and somatosensory cortices, however this was much more widespread than tool identity decoding. We also observed clusters of significant decoding in left SMA, left IPS, left PMd and SPOC bilaterally. More ventral regions exhibiting significant decoding included bilateral LOTC, extending to the left MTG, bilateral para-hippocampal gyrus and right FusiG and AG. Successful decoding of tool-use action in the pantomime task was also observed in the right cerebellum.

### Whole-brain cross-task searchlight

No clusters survived correction in either direction of the cross-task searchlight analysis.

### Spatial overlap of viewing/pantomime searchlight maps

Despite the absence of reliable cross-task decoding, the searchlight maps showed spatial overlap between regions exhibiting significant within-task tool-use action decoding during viewing and pantomime in left MC, bilateral SC, ventral and ventro-dorsal regions including LOTC, fusiform gyrus, and MTG, and dorsal-dorsal regions including IPS and SPOC. Thus, although action-specific information was present in spatially overlapping regions across the two tasks, the absence of cross-task decoding provided no evidence that this information was encoded in a shared format.

## Discussion

Viewing objects can evoke information about their associated actions even when no action is required. Here, we asked how specifically the brain represents these learned object-action associations and whether action information inferred from viewing tools shares a common format with that engaged during tool-use pantomime. Tool-use action could be decoded during viewing across occipitotemporal, parietal, and, remarkably, somatosensory cortex (SC). During pantomime, action information was widespread across occipitotemporal, frontoparietal, and sensorimotor regions. LOTC-hand was particularly notable: it was the only ROI in which tool-use action decoding exceeded within-action tool-identity decoding during both viewing and pantomime. Yet, despite robust within-task decoding and substantial anatomical overlap between tasks, we found no reliable cross-task decoding in either ROI or whole-brain searchlight analyses. Thus, distributed activity patterns contain information about the specific action associated with a viewed tool, but spatial overlap in action information across perception and action did not imply a shared representational format.

A central finding was that specific tool-use actions could be decoded from frontoparietal regions while participants simply viewed tool pictures. Action information was present in IPS and SMG despite participants neither moving their hands nor being instructed to consider how the tools were used. These effects cannot readily be reduced to grip differences because action decoding was matched for grip type. Our findings therefore extend evidence that viewing manipulable objects engages regions implicated in action (Chao and Martin, 2000; Lewis, 2006) by demonstrating that these responses contain information about which action is associated with a viewed tool. SMG may integrate functional and action-related tool knowledge (Binkofski and Buxbaum, 2013; Garcea and Buxbaum, 2019), whereas pMTG contained action information during both viewing and pantomime, consistent with its proposed role in representing learned object-associated actions (Valyear and Culham, 2010; Wurm and Lingnau, 2015). Moreover, action decoding in IPS and PMd during pantomime demonstrates that dorsal regions can represent learned tool-use actions without online visual feedback from the hand or tool.

The presence of tool-use action information in SC during viewing is particularly striking. Although SC is conventionally associated with bodily sensation, its activity can contain information about objects that are only seen: familiar object categories and visually perceived texture can be decoded without haptic stimulation (Smith and Goodale, 2015; Sun et al., 2016). Here, SC discriminated whether a viewed tool would normally be squeezed or rotated. One possibility is that visual recognition of familiar tools reinstates learned somatosensory information associated with their use, potentially including predictions about the sensory consequences of corresponding actions. Consistent with this interpretation, SC contains information about upcoming movements before execution (Gale et al., 2021). Thus, SC may participate not only in processing incoming bodily signals but also in reinstating sensory information associated with familiar actions. A parallel occurs in the gustatory system, where viewing food pictures elicits taste-quality-specific information in gustatory cortex without taste delivery (Avery et al., 2021). Learned associations may therefore allow visual objects to reinstate information within cortical systems associated with the sensory consequences of interacting with them. More broadly, this is compatible with embodied accounts in which object knowledge includes learned bodily and sensory consequences of interaction: recognising a tool may partially reinstate not only what can be done with it, but what performing that action would feel like.

Action information was also prominent in LOTC. Previous work has shown that LOTC contributes to explicit discrimination of actions associated with viewed tools (Perini et al., 2014). Our findings extend this work by showing that specific tool-use actions can be decoded from LOTC when participants simply view tool pictures and action is task-irrelevant. Both LOTC-tool and LOTC-hand contained action information during viewing and pantomime, consistent with evidence that LOTC representations extend beyond visual object form to action-related properties (Bracci and Op de Beeck, 2016; Gallivan et al., 2016; Wurm and Caramazza, 2022). Particularly informative was LOTC-hand, the only ROI in which tool-use action decoding exceeded matched tool-identity decoding in both tasks. This complements our previous finding that LOTC-hand, rather than LOTC-tool, represents how tools should be grasped for use (Knights et al., 2021). Together, these findings are difficult to reconcile with an account of LOTC-hand based solely on visual hand form. Instead, LOTC-hand appears sensitive to action properties linking hands with objects, including how a tool should be grasped and how the hand should move to use it. Thus, the functional organisation of high-level visual cortex may reflect not only what entities look like, but also learned relationships between objects and the bodily actions through which we interact with them.

These findings also speak to the broader concept of affordances - the opportunities for action that objects provide (Gibson, 1979). Whereas our previous work showed that LOTC-hand represents how tools should be grasped (Knights et al., 2021), the present findings extend this relationship to learned tool-use actions. Because grip type was matched across action categories, the present effects cannot readily be reduced to differences in grasp affordance. Rather, they suggest that hand-selective LOTC is sensitive to learned relationships between object perception and characteristic hand actions. However, greater action than tool-identity decoding should not be interpreted as demonstrating an identity-independent action code; rather, it indicates greater sensitivity to associated action categories than to the matched within-action identity distinctions tested here. Future cross-identity generalisation studies could determine whether such action information generalises across visually distinct tools. Tool-use information in LOTC during pantomime, when neither the tool nor hand was visible, further suggests that these responses do not depend on concurrent visual hand or object input. This accords with action-related LOTC responses without visual input (Astafiev et al., 2004; Gallivan et al., 2016) and tool-related responses in congenitally blind individuals (Peelen et al., 2013). LOTC may therefore form part of a distributed system linking perceptual object information with learned knowledge about body-object interactions.

Nevertheless, the distribution of action information was strongly task-dependent. During viewing, decoding was generally stronger in occipitotemporal regions, whereas during pantomime it was stronger throughout frontoparietal and sensorimotor cortex. This is compatible with distinctions between ventral and dorsal contributions to perception and action (Milner and Goodale, 2006) but argues against strict separation: action information was present in dorsal and somatosensory regions during perception, whereas ventral regions represented action information during production. Although our individually defined ROIs focused on the left hemisphere based on the established left lateralization of tool processing, whole-brain searchlight decoding found substantially bilateral effects. Thus, left-hemisphere specialization for tool processing does not imply that action information is confined to the left hemisphere.

Critically, our findings directly address whether action information elicited during tool viewing shares a common format with production of the corresponding action. We found no reliable cross-task decoding of tool-use action in any ROI or the whole-brain searchlight. Thus, overlapping activation, or successful decoding of the same property in two tasks, does not establish that the underlying multivoxel patterns are shared. Our findings are consistent with action-related information being represented in task-dependent formats according to whether it is inferred from visual input or recruited during action production. Comparable dissociations occur in other domains: inferred and experienced tastes can be decoded from gustatory cortex without sharing reliably generalisable activity patterns (Avery et al., 2021), while action information during motor imagery and action planning can occur in visual cortex without corresponding cross-task generalisation (Monaco et al., 2020). Representation of the same information within a cortical region therefore need not imply a task-invariant neural code. More broadly, perception-action coupling associated with familiar tools need not depend on reinstating the same neural activity patterns used to produce the corresponding actions. Instead, our findings are compatible with action information being transformed into task-appropriate representational formats as the system moves from perceptual inference towards action production.

This distinction has broader relevance for computational accounts of visually guided action. Both biological and artificial agents face the problem of transforming information about object properties into appropriate actions. Our findings suggest that biological perception-action coupling need not rely on a single representation carried unchanged from object recognition to action production. Instead, information about the same action may be expressed differently as perceptual knowledge is transformed into behaviour. This provides a potential biological constraint for computational and robotic models seeking to connect object recognition with flexible action, although determining the transformations involved will require direct investigation.

The lack of significant cross-decoding should nevertheless be interpreted cautiously. The tasks differed not only in perceptual versus motor demands but also in their immediate inputs: viewing was elicited by tool pictures, whereas pantomime was cued by tool names. Although cross-format decoding between pictures and words has been demonstrated for semantic object information (Fairhall and Caramazza, 2013; Shinkareva et al., 2011), differences in stimulus format may have reduced sensitivity to shared representations. Moreover, Chen et al. (2018) reported cross-task decoding of tool-use information in left inferior parietal cortex, but their perceptual task explicitly encouraged consideration of object features, including associated actions, whereas our participants performed a one-back identity task. Cross-task invariance may therefore depend on region, abstraction level, and cognitive context rather than being an obligatory property of tool representations. Explicit attention to or imagery of a tool’s associated action may generate a more production-like neural state than action information elicited when action is task-irrelevant.

Pantomime also differs importantly from real tool use. It requires retrieving and producing a learned action without the physical constraints, visual feedback, or somatosensory consequences of interacting with an actual tool. Real and pantomimed tool use differ in activation, multivoxel patterns, and functional connectivity (Chen et al., 2023), while neural and behavioural responses to real objects can differ from pictures (Snow and Culham, 2021), and apraxic performance can improve with real objects compared with pantomime (Goldenberg et al., 2004). Thus, failure to cross-decode should not imply that perceptual and action representations can never share a common format. Future work should determine whether viewing and subsequently acting upon real, graspable tools produces greater representational correspondence and clarify how perceptual information is transformed into action during genuine object interaction.

In conclusion, simply seeing a tool provides the brain with specific information about how that object is used. Tool-use action could be decoded throughout occipitotemporal and parietal cortex and even from SC in the absence of movement. LOTC-hand showed greater sensitivity to tool-use action than to matched tool-identity distinctions during both viewing and pantomime, supporting its role in learned hand-object interactions. Yet, we found no evidence that action representations were invariant across viewing and pantomimed action production. Together, our findings suggest that perception can evoke specific action and sensory information across distributed cortical networks without necessarily recreating the neural state associated with performing or experiencing it. Skilled object interaction may therefore depend not on simple reinstatement of a single perception-action representation, but on the transformation of learned information about objects, bodily actions, and their sensory consequences according to current behavioural demands.

## Authors’ Contribution Statement

**Annie Warman**: Data curation; Formal analysis (lead); Software; Validation; Visualization; Writing - original draft preparation. **Diana Tonin**: Data curation; Investigation (lead); Methodology; Formal analysis; Writing - review & editing. **Fraser W. Smith**: Conceptualization; Methodology; Software (lead); Formal analysis; Writing - review & editing. **Stéphanie Rossit**: Conceptualization (lead); Data curation; Formal analysis; Funding acquisition; Investigation; Methodology; Project administration; Resources; Supervision; Validation; Visualization; Writing - original draft preparation; Writing - review & editing.

## Acknowledgements

We would like to thank the staff at the Scannexus Imaging Center (Maastricht, The Netherlands) for support in data collection and Dr. Stefania Bracci for sharing the visual localizer stimuli. This work was funded by the University of East Anglia.

