## Supplemental text for "Human see, human do? Viewing tool pictures evokes tool-use action information in hand-selective occipitotemporal cortex"

**Supplementary Materials**

**Univariate analysis.** To test whether decoding effects could be explained by differences in mean response amplitude, we extracted β weights for each tool identity, task, and ROI and calculated univariate difference scores corresponding to the comparisons used in the MVPA.

For individual tool identity, β-weight differences were calculated between tool pairs matched for tool-use action and grip type (tongs-nutcracker, key-screw, screwdriver-corkscrew, and tweezers-peg) and then averaged across pairs. For tool-use action, β weights were first averaged for tools sharing the same action and grip type (key/screw, screwdriver/corkscrew, peg/tweezers, and tongs/nutcracker). Differences between rotate and squeeze tools were then calculated separately for precision- and power-grip tools and averaged across grip types. Thus, the univariate comparisons were matched to those used for the MVPA.

β-weight difference scores were entered into a 12 (ROI) × 2 (property: individual tool identity, tool-use action) × 2 (task: viewing, pantomime) repeated-measures ANOVA. Post-hoc comparisons were FDR corrected for multiple comparisons (Benjamini & Yekutieli, 2001).
